# Human Plasma-Like Medium (HPLM) induces *Cryptococcus neoformans in vivo* cell morphologies

**DOI:** 10.1101/2023.08.29.555343

**Authors:** Eduardo G Mozo, Orlando Ross, Raif Yuecel, Ivy M. Dambuza, Liliane Mukaremera

**Affiliations:** Medical Research Council Centre for Medical Mycology at The University of Exeter, University of Exeter, Exeter, United Kingdom; Exeter Centre for Cytomics, The Bioeconomy Centre, Faculty of Health and Life Sciences, University of Exeter, Exeter, UK

**Keywords:** *Cryptococcus neoformans*, cell morphology, HPLM, titan cells

## Abstract

The fungal pathogen *Cryptococcus neoformans (C. neoformans)* forms yeast cells of different sizes and morphological characteristics during infection. These features are usually not seen in standard laboratory *in vitro* conditions. Here, we describe *in vivo* cell morphologies when *C. neoformans* is grown in human plasma-like medium at 37°C-5% CO_2_. We observed mixed-size populations of cells less than 1 μm up to 16.8 μm cell diameter, increased capsule size, high chitin and DNA content in larger cells. Our findings show serum is not required for HPLM-induced *C. neoformans* cellular heterogeneity. Thus, this new method offers an opportunity to investigate factors of *C. neoformans* that mediate pathogenesis or host-pathogen interactions in a physiologically relevant setting.

**IMPORTANCE:** Description of new *in vitro* culture condition using the human plasma-like medium that supports the formation of the full range of *in vivo* cell morphologies of *C. neoformans*.

---

The fungal pathogen *C. neoformans* forms yeast cells of different sizes and morphological characteristics *in vivo*. Some of these dynamic cell populations include typical sized yeasts (5-7 μm), enlarged cells referred to as titan cells (> 10 μm), and small cells (less than 5 μm) that include micro cells, drop cells, seed cells and titanides (1-4). In contrast to *in vivo* conditions where significant size heterogeneity exists, *C. neoformans* grown under nutrient rich conditions *in vitro* predominantly produce typical sized yeast cells with a uniform morphology. Most *in vitro* studies on *C. neoformans* have used these homogenous cell populations, which although convenient do not represent the broad range of cellular morphologies present during infection, and therefore may lead to inaccurate conclusions about factors that mediate *C. neoformans* host-pathogen interactions and pathogenesis.

In 2018, three protocols for the production of titan cells *in vitro* were described (4-6). The factors found to induce the formation of titan cells include: (a) low nutrients, hypoxia, low pH, and continuous shaking (5), (b) low nutrients, neutral pH, presence of serum and azide in static conditions with a CO_2_ enriched atmosphere (6), and (c) low nutrients and presence of serum in static conditions with a CO_2_ enriched atmosphere (4). Although there are differences between these three protocols, there are also common themes, such as growth under low nutrient, oxygen limitation and low cell density conditions.

Here we describe new *in vitro* conditions that induce the formation of cell morphologies that mimics the diversity of cell morphologies observed *in vivo*. To induce the formation of *in vivo* like *Cryptococcus* cell morphologies, the wild type H99 strain was used. Overnight cultures in nutrient-rich YPD were inoculated into the “human plasma-like medium” (HPLM) and then incubated at 37°C-5% CO_2_. HPLM is a relatively new culture medium that contains polar metabolites similar to those present in the human plasma (7). We observed that *Cryptococcus* can grow and multiply in HPLM medium. Microscopic observation of cells grown in HPLM show a population of cells with sizes ranging from small cells of less than 1 μm to large cells up to 16.8 μm in cell diameter (Fig 1A-C). Using previously described cell body size (diameter) measurements, C. neoformans cells were divided into 3 sub-populations; titan cells (>10 μm), normal-sized yeasts (5-9 μm) and smaller cells (<4 μm). We observed a mixture of all 3 cell sub-populations in our cultures (Fig 1A-C). Similar to previous observations, low cell density induced more cells with a diameter of more than 10 microns when compared to high cell density inocula at 48 and 168 h of incubation (Fig 1B and Fig 1C, respectively). We also tested strains known to have defects in titan cell formation *in vivo* and *in vitro*, the *rim101*Δ and *gpr4*Δ/*gpr5*Δ mutants (8). Our results show that these two mutants had defects in large cell formation (>10 microns) in HPLM (Fig 1D). *In vivo* titan cells are polyploid and have an increased chitin content. Therefore, we used flow cytometry to examine the DNA and cell wall chitin content after staining with propidium iodide (PI) and Calcofluor White (CFW), respectively. We analysed *Cryptococcus* grown in HPLM and nutrient rich YPD (control). Typical 5-7 μm cells in YPD had 1C and 2C DNA contents (population 1, Fig 1E). Similar sized cells (population 1) grown in HPLM grown cells also had 1C and 2C DNA content. Notably, a smaller size population of cells was also observed that had varying side scatter properties, suggestive of cellular diversity (population 6, Fig 1E). This unique population observed in HPLM was predominantly comprised of cells with 1C DNA content. Consistent with published reports (5, 9), the large cells (populations 4 and 5) had higher DNA content that was >2C (Fig 1E). Similarly, the large cells (> 10 μm) had increased cell wall chitin compared to the typical sized and small cells in HPLM (Fig 1F-G) as previously described for *in vivo* titan cells (5, 10).

**Figure 1.**
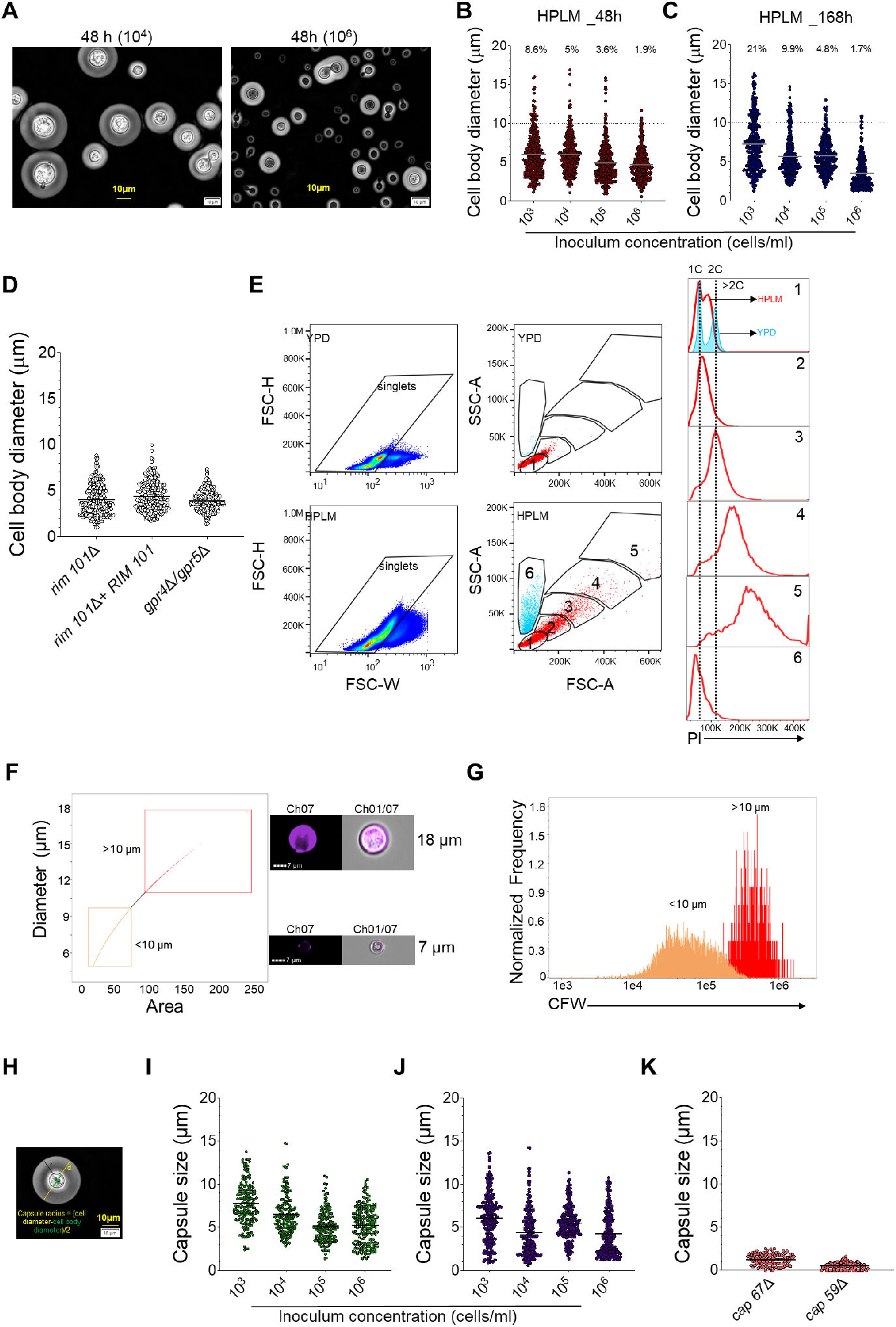
*Cryptococcus neoformans* forms *in vivo*-like morphologies in HPLM at 37°C-5% CO2. *C. neoformans* H99 and various mutant strains were grown overnight in YPD at 30°C with shaking (200 rpm), then washed twice with sterile water and counted with haemocytometer. Various inoculum concentrations were added to 6-well plates containing HPLM media, and incubated at 37°C, 5% CO_2_ for 48 h or 168 h. After incubation, *C. neoformans* cells were analysed for their cell body (diameter), capsule sizes, DNA content and chitin content. **A**, *C. neoformans* cells were fixed with formaldehyde, suspended in India ink and imaged on an Olympus CKX53 microscope. **B-C**, Cell diameters were measured using ImageJ at 48h (B) and 168h (C). Data presented are representative of 3 biological replicates with at least 300 cells. **D**, Cell size of mutants with defects in titan cell formation. 10^4^ cells/ml were cultured in HPLM medium as described above. Data presented are representative of 4 biological replicates with at least 300 cells measured. **E**, *C. neoformans* H99 cells grown in YPD or HPLM for 48 h, washed in cold sterile water, fixed with 80% methanol and stained with 50 µg/ml PI for 30 min. DNA content was measured using flow cytometry for populations 1-6. **F-G**, To measure chitin content, 10^6^ cells/ml were stained with 25 μg/ml Calcofluor White for 5 minutes, washed and then analysed by imaging flow cytometry. Calcofluor White (for chitin content) stained cells were analysed by imaging Flow Cytometer. Data presented are from 8,544 singlets. **H**, Diagram of capsule radius measurement. **I-J**, Capsule thickness/radius of *C. neoformans* H99 after 48h (I) and 168 h (J) of incubation in HPLM. Data presented are representative of 4 biological replicates with at least 160 cells measured. **K**, Capsule thickness of mutants with defects in capsule formation. 10^4^ cells/ml were incubated in HPLM and data presented are representative of 3 biological replicates with at least 100 cells measured. The grey line in figures represents the median. The dotted line at Y-axis represents the 10-micron cut-off, and the numbers the percentage of cells above 10-micron DNA: Deoxyribonucleic acid, PI: propidium iodide, YPD: yeast extract-peptone-dextrose.

In addition to various cell sizes, cells grown in HPLM media had a large capsule (Fig 1A and H). In HPLM, the capsule radius varied between 1.6 to 13.5 μm (median 6.3) in titan cells and 0.6 to 12.8 μm (median 5.4) in non-titan cells (Fig 1I-J). This is bigger than previously described *in vitro* cells that had a median capsule radius of 4.8 μm in titan and 2.7 μm in typical cells *in vitro* (5). Interestingly, some HPLM-grown cells displayed capsule sizes up to 13.5 μm, which is close to sizes observed for *in vivo* titan cells (14.8 μm) (5). Mutants with known defects in capsule formation (*cap59*Δ and *cap67*Δ) (11, 12) showed small to no capsule formation in HPLM (Fig 1K). These findings show that the large cells formed in HPLM possess the characteristics of *in vivo* titan cells.

Small *C. neoformans* cells were previously characterised depending on various morphological factors, and have been designated by multiples names including micro cells (∼1μm) (13), titanides (oval, metabolically active with a thin cell wall) (4), drop cells (metabolically inactive with a thick cell wall) (14), or seed cells (similar to titanides but have increased mannose exposure and are seen in cultures devoid of titan cells) (3). Based on the varying side scatter properties of these small cells, we hypothesise that the small cell population generated in HPLM contains a mixture of small cell morphologies.

Serum has been previously described to be an inducer of titan cell formation *in vitro* (4, 6). We supplemented the HPLM medium with serum to determine whether it would enhance the production of titan cells. *C. neoformans* grew slowly in HLPM alone (Fig 2A) contrary to the fast growth observed in HPLM supplemented with 10% fetal bovine serum (FBS) (Fig 2B). There was also an increase in the number of titan cells in the presence of FBS, both at 48 h (Fig 2C) and 168 h of incubation (Fig 2.D). For example, titan cells (>10 μm) represented 35.4% of the whole population at 168 h in the presence of serum, while they were 21% in HPLM alone when the initial inoculum was 10^3^ cells (Fig 1C and Fig 2D). DNA content was similar in both HPLM and HPLM supplemented with 10% FBS (Fig 2F, overlaid on HLPM alone). In addition, capsule sizes were comparable in HLPM or HPLM supplemented with serum (Fig 2H-J). These data show that the presence of serum provided a boost in *C. neoformans* titan cell formation, but HPLM alone was sufficient to induce cellular heterogeneity similar to that observed in *in vivo*.

**Figure 2.**
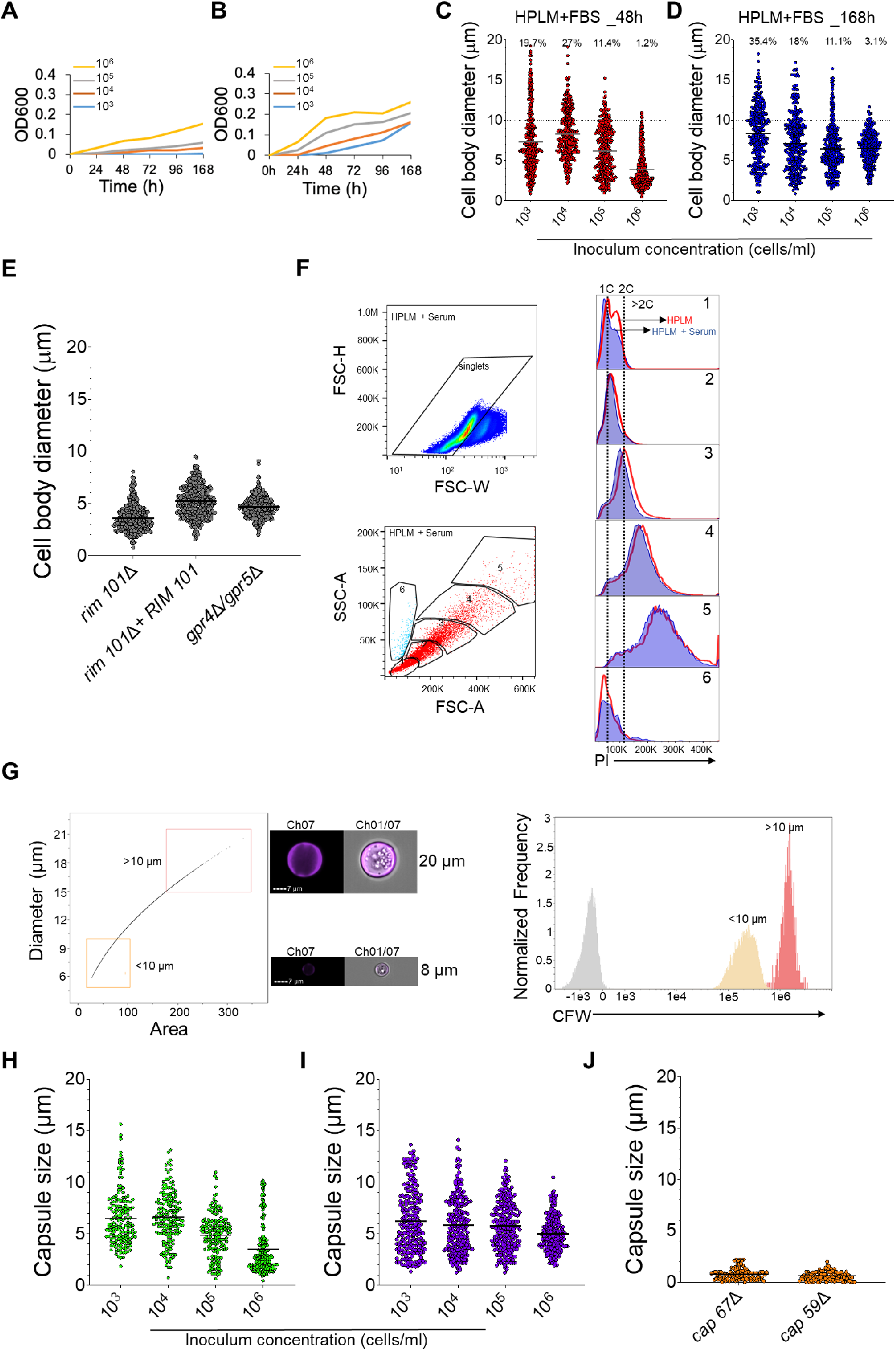
Serum is dispensable for HPLM-induced *Cryptococcus neoformans* cellular heterogeneity. *C. neoformans* H99 and various mutant strains were grown overnight in YPD at 30°C with shaking (200rpm), then washed twice with sterile water and counted with hemocytometer. Various inoculum concentrations were added to 6 well plates containing HPLM medium supplemented with 10% FBS, and incubated at 37°C, 5% CO_2_ for 48 h or 168 h. After incubation, *C. neoformans* cells were analysed for their cell body (diameter), capsule sizes, DNA content and chitin content. **A-B**, Growth curve of *C. neoformans* H99 in HPLM (A) and HPLM supplemented with FBS (B). Optical density (OD600) was read at 0, 24, 48, 72, 96 and 168 h of incubation. **C-D**, Cell body sizes of *C. neoformans* H99 in HPLM supplemented with FBS at 48 h (C) and 16 8h (D) of incubation. Data presented are representative of 3 biological replicates with at least 300 cells. **E**, Cell body sizes of mutants with defects in titan cell formation at 48 h of incubation. Data presented are representative of 4 biological replicates with at least 300 cells measured. **F**, DNA content of H99 cells grown in HPLM (red line, overlaid Fig 1E) and HPLM supplemented with 10% FBS (blue background). **G**, To measure chitin content, 10^6^cells/ml were stained with 25 μg/ml Calcofluor White for 5 minutes, washed and then analysed by imaging flow cytometry. **H-I**, Capsule thickness (radius) of H99 cells grown in HPLM and 10%FBS at 48 h (H) and 168 h of incubation (I). Data presented are representative of 4 biological replicates with at least 160 cells measured. **J**, Capsule thickness of mutants with defect in the capsule formation grown in HPLM supplemented with FBS at 48 h incubation. Data presented are representative of 3 biological replicates with at least 100 cells measured. The grey line in figures represents the median. The dotted line at Y-axis represents the 10-micron cut-off, and the numbers the percentage of cells above 10-micron. DNA: Deoxyribonucleic acid, FBS: Foetal Bovine serum, PI: propidium iodide.

Collectively, the results presented here show that HPLM, a medium that more closely resembles the human plasma, incubated at 37°C with 5% CO_2_ can be used to induce the diversity of *C. neoformans* cell morphologies observed *in vivo*. HPLM has been used for immunological studies where it induced a different transcriptional response in human primary T-cells and improved their activation after antigen stimulation (15). To our knowledge, HPLM has not been used to grow fungal pathogens and we show for the first time that *C. neoformans* grows and differentiates into *in vivo* cell morphologies in this medium. Thus, HPLM is a great option to use in experiments investigating *C. neoformans* pathogenesis, specifically host-*Cryptococcus* interactions, as it can be used to culture both the fungus and the host immune cells. For example, HPLM is the optimal growth medium when co-culturing *C. neoformans* with human primary T-cells *in vitro*. Future work should focus on identifying how this media influences *Cryptococcus* gene expression that leads to the formation of different cell morphologies in *C. neoformans* and other members of the *Cryptococcus* species complex.

## ACKNOWLEDGMENTS

We thank Prof Kirsten Nielsen for providing us with *C. neoformans* strains used in this study. We also thank Prof Neil Gow, Ms Marina Yoder and Prof Kirsten Nielsen for helpful feedback on the manuscript.

LM is supported by the Academy of Medical Sciences/the Wellcome Trust/ the Government Department of Business, Energy and Industrial Strategy/the British Heart Foundation/Diabetes UK/Global Challenges Research Fund Springboard Award [SBF006\1142]. OR is supported by the MRC Doctoral training Grant MR/W502649/1. IMD is supported by funding from the Wellcome Trust (102705, 217163). EDM, OR, IMD and LM are also supported by the Medical Research Council Centre for Medical Mycology at The University of Exeter (MR/N006364/2 and MR/V033417/1).

LM and IMD conceived the study, performed most experiments, interpreted data and wrote the manuscript. EGM, OR and RF performed experiments and analysed the data. LM, IMD, and RF wrote and edited the manuscript.

